# A Novel Drosophila Model of Cancer Cachexia Reveals Conserved Jak/Stat Dependent Glucagon-like Akh Activation as a Driver of Metabolic Reprogramming and Systemic Wasting

**DOI:** 10.1101/2025.09.10.675448

**Authors:** Kewei Yu, Gurpreet S. Moroak, Esther M. Verheyen

## Abstract

Cancer-associated cachexia is a systemic wasting syndrome with no effective therapies, and results in millions of deaths annually. Here, we establish a novel *Drosophila* model of cancer cachexia using overexpression of Hipk and constitutively active Sik3 in larval epithelial tissue. Tumor-bearing larvae has significant muscle and fat body wasting, together with elevated carbohydrates and lipolysis. Mechanistically, tumors secrete Unpaired ligands that activate Jak/Stat signaling in corpora cardiaca (cc) cells, inducing the expression and protein levels of glucagon-like hormone Adipokinetic hormone (Akh). Elevated Akh, together with the lipase Brummer (Bmm), drives the aforementioned systemic metabolic reprogramming and tissue catabolism. In conclusion, this study identifies a conserved tumor-host Upd–JAK/STAT–Akh signaling axis that contributes to organ wasting.

## Introduction

Cancer-associated cachexia is a systemic paraneoplastic syndrome characterised by substantial weight loss due to loss of skeletal and adipose tissue and is estimated to cause an annual death rate of 2 million people worldwide (Baracos et al., 2018; Farkas et al., 2013; Fearon et al., 2011; Ferrer et al., 2023). These alterations in distant organs away from the site of the tumor are a result of the tumor altering the host’s systemic metabolism (Fearon et al., 2011; Porporato, 2016; Swanton et al., 2024). Thus far, there is no Food and Drug Administration (FDA) approved drug therapy that can treat or reduce the progression of cancer cachexia (Baracos et al., 2018). This highlights the need to use model organisms to investigate the underlying mechanisms of tumour-host interactions resulting in cancer cachexia to facilitate potential new therapies targeting factors promoting cancer cachexia.

Drosophila is a well-established model for investigating cancer, with highly conserved genes and cell signaling pathways, as well as physiological conservation of organ functions (Bilder et al., 2021). In the recent decade, Drosophila tumor model studies have elucidated numerous cachexic factors and underlying signalling pathways mediating systemic organ wasting (Liu et al., 2022). For example, cachexic factors such as interleukin/unpaired (Upd) ligands, insulin-like growth factor binding protein (IGFBP)/ImpL2, VEGF/Pvf1 and bone morphogenetic protein (BMP)/Gbb have been found to induce host organ wasting coupled with abnormal metabolite levels in several different Drosophila adult and larval tumor models (Ding et al., 2021; Figueroa-Clarevega & Bilder, 2015; Kwon et al., 2015; Lodge et al., 2021; Song et al., 2019). Studies of human cancer cachexia patients have found that serum IGFBP2 is a biomarker for muscle wasting in pancreatic ductal adenocarcinoma patients (Dong et al., 2021) and interleukin-6 has been found to promote cancer cachexia in multiple types of human cancer and mouse models of cancer cachexia (Agca & Kir, 2024). These parallels show that Drosophila is a powerful model for understanding potential mechanisms of cancer cachexia.

Homeodomain-interacting protein kinases (Hipks) regulate cell proliferation, apoptosis, and tissue development (Blaquiere & Verheyen, 2017). Previously, our lab established a larval tumor model based on overexpression of Hipk in Drosophila epithelial tissue. In this model, we observed hallmarks of cancer such as neoplasia, epithelial to mesenchymal transition (EMT) and increased aerobic glycolysis (Blaquiere et al., 2018; Wong et al., 2019). More recently we described a tumour synergy between Hipk and Salt-inducible kinases (Sik2 and Sik3) (Yu et al., 2023). Siks are serine/threonine kinases in the AMPK family (Wein et al., 2018). Humans possess 3 SIKs while the fly genome encodes Sik2 and Sik3 (Choi et al., 2011). Siks can integrate conserved signaling pathways like Notch and Hippo to promote tumor growth in Drosophila (Şahin et al., 2020; Wehr et al., 2013). Co-expression of constitutively active forms of Sik2 or Sik3 (Sik3-CA) with Hipk caused significant tissue hyperplasia and tissue distortion, accompanied by elevated dMyc, Armadillo/β-catenin, and the Yorkie target gene expanded indicating that both Sik2 and Sik3 can synergize with Hipk to promote tumorous phenotypes (Yu et al., 2023). The human homolog of *dMyc*, *MYC* is a well-characterized proto-oncogene and is estimated to have elevated or deregulated expression in up to 70% of human cancers (Dang, 2012). Wnt/β-catenin and the Hippo signaling pathway are also both often deregulated in human cancers (Harvey et al., 2013; Zhang & Wang, 2020). These findings indicate that Hipk and Sikare able to promote tumorigenesis through these conserved signaling pathways and through the proto-oncogene Myc.

In the current study, we establish the ‘Hipk+Sik’ Drosophila tumor model as a novel cancer cachexia model by showing that hyper- and neoplasia induced by overexpression of Hipk and Sik3-CA in larval epithelial tissue causes muscle and fat wasting away from the site of the tumor. We characterized the aberrant metabolism in these tumor-bearing larvae and observed increased carbohydrates and lipolysis (breakdown of triacylglycerol). We determined that these metabolic reprogramming is induced by both glucagon-like adipokinetic hormone (Akh) and Adipose triglyceride lipase (ATGL)/Brummer (Bmm). Finally, we show that Unpaired ligands secreted from the tumor tissue activate JAK/STAT signalling in corpora cardiaca (cc) cells to induce Akh expression and protein levels. Therefore, we propose a Drosophila cancer cachexia model whereby tumors secrete Unpaired ligands to induce Akh in the distant organ cc cells, and Akh and Bmm converge to contribute to the aberrant metabolism, contributing to distant organ wasting.

## Results

### *dpp>Hipk+Sik3-CA* induces functional changes in muscle and muscle wasting in Drosophila larvae

Ectopic expression of Sik3-CA and Hipk in imaginal discs using the *dpp-Gal4* driver strain *(dpp>Hipk+Sik3-CA)* induces a prolonged third instar larval phase and delayed pupariation (Yu et al., 2023). Progeny expressing Hipk and Sik3-CA undergo progressive size growth over time and around two-thirds are unable to pupariate and die as giant larvae (Fig. 1A, Fig. S1A-B). To investigate this phenomenon, we chose 2 different developmental timepoints for the tumor model larvae. As overexpression of *Hipk* and *Sik3-CA* induces a developmental delay, we used the 50% pupariation rate to developmentally stage match with the control *dpp>GFP + RFP* larvae (Yu et al., 2023). This is to ensure that the experimental group is at the same life stage as the control for consistent comparisons. Thus, we consider D5 control larvae age matched with D7-8 *dpp>Hipk+Sik3-CA* ‘Tumor’ larvae. ‘Cachexic’ larvae are defined as D16 *dpp>Hipk+Sik3-CA* larvae which are bloated (significant increase in extractable hemolymph volume) and significantly bigger in mass (Fig. 1A-C). Such a tumor induced bloating phenotype has been shown since the 1960s and across both larval and adult Drosophila tumor models (Brumby, 2003; Gateff & Schneiderman, 1969; Pagliarini & Xu, 2003). Numerous Drosophila tumor models with this bloating phenotype are also characterized as cancer cachexia models (Figueroa-Clarevega & Bilder, 2015; Khezri et al., 2021; Santabárbara-Ruiz & Léopold, 2021). Therefore, we hypothesize that the increase in larval mass and transparency of the larval body indicative of loss of fat body indicates that our *dpp>Hipk+Sik3-CA* tumor model could also induce distant organ wasting reminiscent of human cancer cachexia-like syndrome.

**Figure 1.**
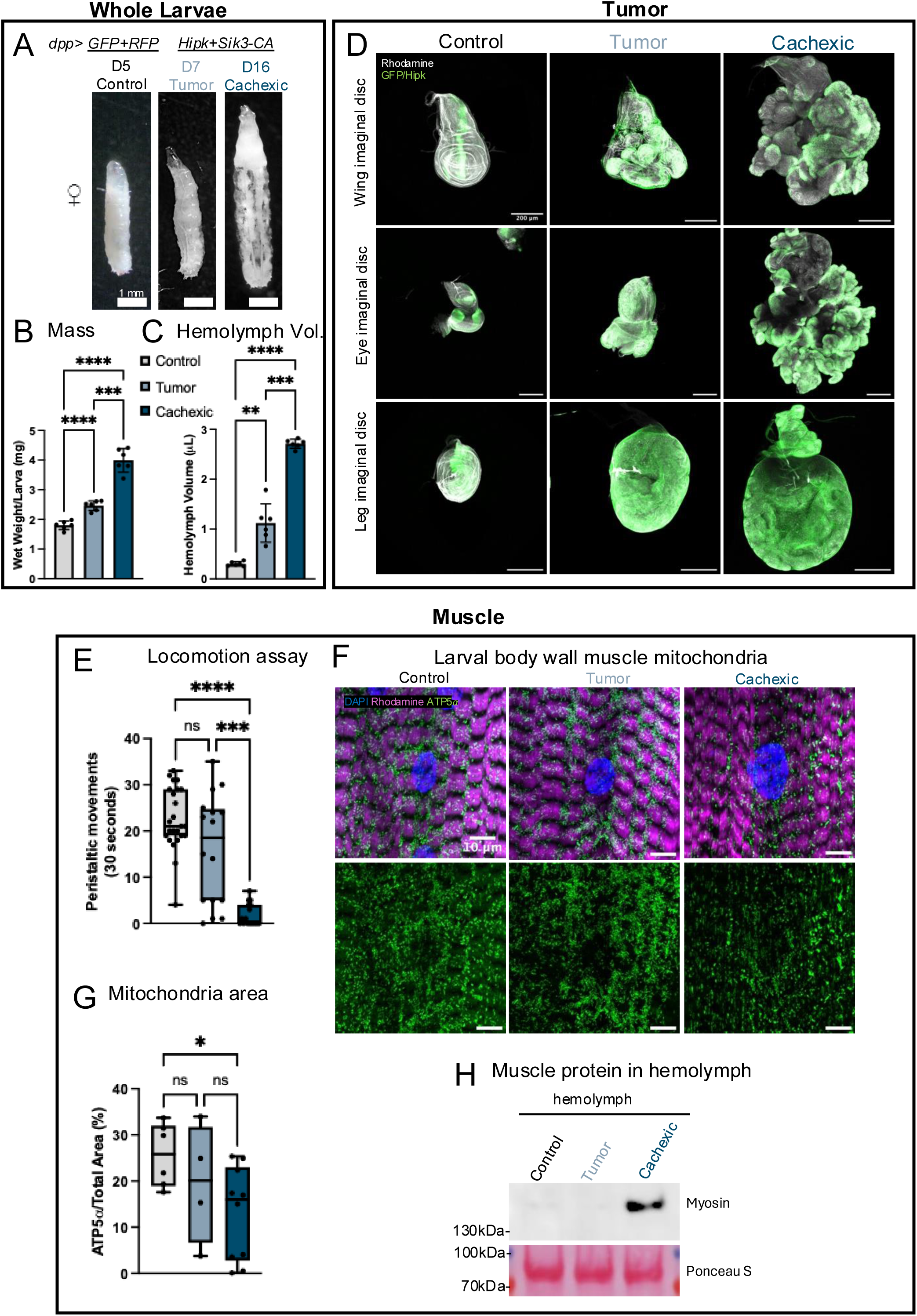
*dpp>Hipk+Sik3-CA* induces functional changes in muscle and muscle wasting in Drosophila larvae. (A) Representative images of control, tumor-bearing and cachexic larvae (*n* = 9, 13 and 15 respectively). (B) Wet weight of larvae with the indicated genotypes (*n*=5/data point) ^∗∗∗^p = 0.0001, and ^∗∗∗^*p < 0.0001 (Brown-Forsythe and Welch ANOVA test). (C) Hemolymph volume in larvae with the indicated genotypes (*n*=5/data point) ^∗∗^p =0.0055, ^∗∗∗^p = 0.0006, ^∗∗∗^*p < 0.0001 (RM one-way ANOVA). (D) Representative maximum projection confocal images of F-actin staining (gray), GFP and Hipk is green (*n*>4 for all discs). (E) Quantification of number of peristaltic movements within 30 seconds (*n*=24, 16 and 17 respectively) ns=0. 1670, ^∗∗∗^p = 0.0006, ^∗∗∗∗^p < 0.0001 (Brown-Forsythe and Welch ANOVA test). (F) Longitudinal maximum projection confocal images of ventral longitudinal muscle VL4 from A2-A4 hemisegments. Anti-ATP5α (green), F-actin (red), and DAPI (blue) (n=7, 4 and 9 respectively). (G) Quantification of area of ATP5α as a percentage of total area of muscle. ns=0.775 and 0.7591 respectively, and ^∗^p = 0.00293 (Brown-Forsythe and Welch ANOVA test). (H) Western blot of whole protein extracts of hemolymph from third instar larvae and probing for the presence of sarcomere protein Myosin. Ponceau S is used as a loading control.

Over the progression of the increased bloating from tumor-bearing to cachexic, we also observe a drastic increase in tumor size and changes in epithelial tissue morphology in all epithelial discs overexpressing Hipk and Sik3-CA in the *dpp-Gal4* domain, suggesting an increase in tumor-burden of the larvae over time (Fig. 1D). In these discs, Hipk immunostaining marks the dpp-Gal4 expression domain and label the tumor cells. In addition, we observed significantly reduced locomotion in cachexic larvae compared to control and tumor-bearing (Fig. 1E) with most cachexic larvae with no peristaltic movements within 30 seconds. Drosophila larvae move forward by peristaltic muscle contractions, and locomotion is a measure of muscle and neuronal function (Weitkunat & Schnorrer, 2014). As muscle mitochondria generates Adenosine Triphosphate (ATP) for peristaltic movement, we immunostained larval muscle to detect ATP synthase F1 Subunit Alpha (ATP5α), a mitochondrial marker. Body wall muscle mitochondrial content in cachexic larvae is significantly reduced compared to control, suggesting that reduced mitochondria might be contributing to reduced muscle contraction and therefore impaired muscle function (Fig. 1F, G).

Due to the significantly reduced muscle function of cachexic larvae, we wondered if this was accompanied by muscle wasting/degeneration as muscle weakening and wasting frequently occur together. While overall body wall muscles did not appear abnormal when staining with rhodamine-conjugated phalloidin (Fig. 1F), we characterized muscle wasting by probing for the major muscle component (Rodríguez-Vázquez et al., 2024). We detected Myosin Heavy Chain (MHC) protein in the hemolymph of cachexic larvae but not control or tumor-bearing larvae (Fig. 1H), suggesting that cachexic larval muscle is releasing MHC into circulation, likely due to muscle wasting.

Together, the reduced muscle mitochondria contributing to reduced locomotion and secretion of muscle protein into the hemolymph suggests that *dpp>Hipk+Sik3-CA* can induce non-autonomous distant muscle wasting.

### *dpp>Hipk+Sik3-CA* induces morphological and functional changes in the fat

In addition to muscle wasting, human cancer cachexia patients also suffer from adipose tissue wasting (Argilés et al., 2014). The larval fat body (considered analogous to the human liver and fat tissues) is composed of a single layer of attached polygonal cells that form a large flat structure occupying the body cavity (Meschi & Delanoue, 2021). As we see increased transparency of cachexic larvae indicative of fat body loss (Fig. 1A), we wanted to characterize the fat body morphology change and functional changes potentially induced by the tumors. We find that cachexic larval fat bodies dissociate into spherical cells that are significantly smaller (Fig 2A-C). This phenotype is reminiscent of what occurs normally during Drosophila pupal metamorphosis, wherein matrix metalloproteinases 1 (MMP1) is induced in the third instar larval fat body and is undetectable 6 hours after pupariation, while the E-Cadherin-mediated cell-cell junctions are cleaved by MMP1, resulting in complete dissociation of fat body into individual cells free-floating in the pupal hemolymph (Jia et al., 2014). Additionally, this fat body cell rounding is also observed in another Drosophila cancer cachexia model, in which tumor-secreted MMP1 disrupts the basement membrane/extracellular matrix proteins at the adipocyte cell-cell adhesion junctions, resulting in rounding of fat cells (Lodge et al., 2021). As expected, there is a significant reduction in Dcad2 protein levels in fat cell-cell junction in cachexic larval fat cells (Fig. S2A-B). Interestingly, while there are sustained levels of MMP1 protein at the fat cell-cell junction (Fig. S2B), transcriptional levels of MMP1 are significantly reduced in tumor-bearing and cachexic larval fat tissue (Fig. S2C), suggesting an alternate tissue source of MMP1. This could potentially be mediated by significantly increased levels of MMP1 protein from the cachexic larvae tumor tissue (Fig. S2D) being secreted into the hemolymph.

**Figure 2.**
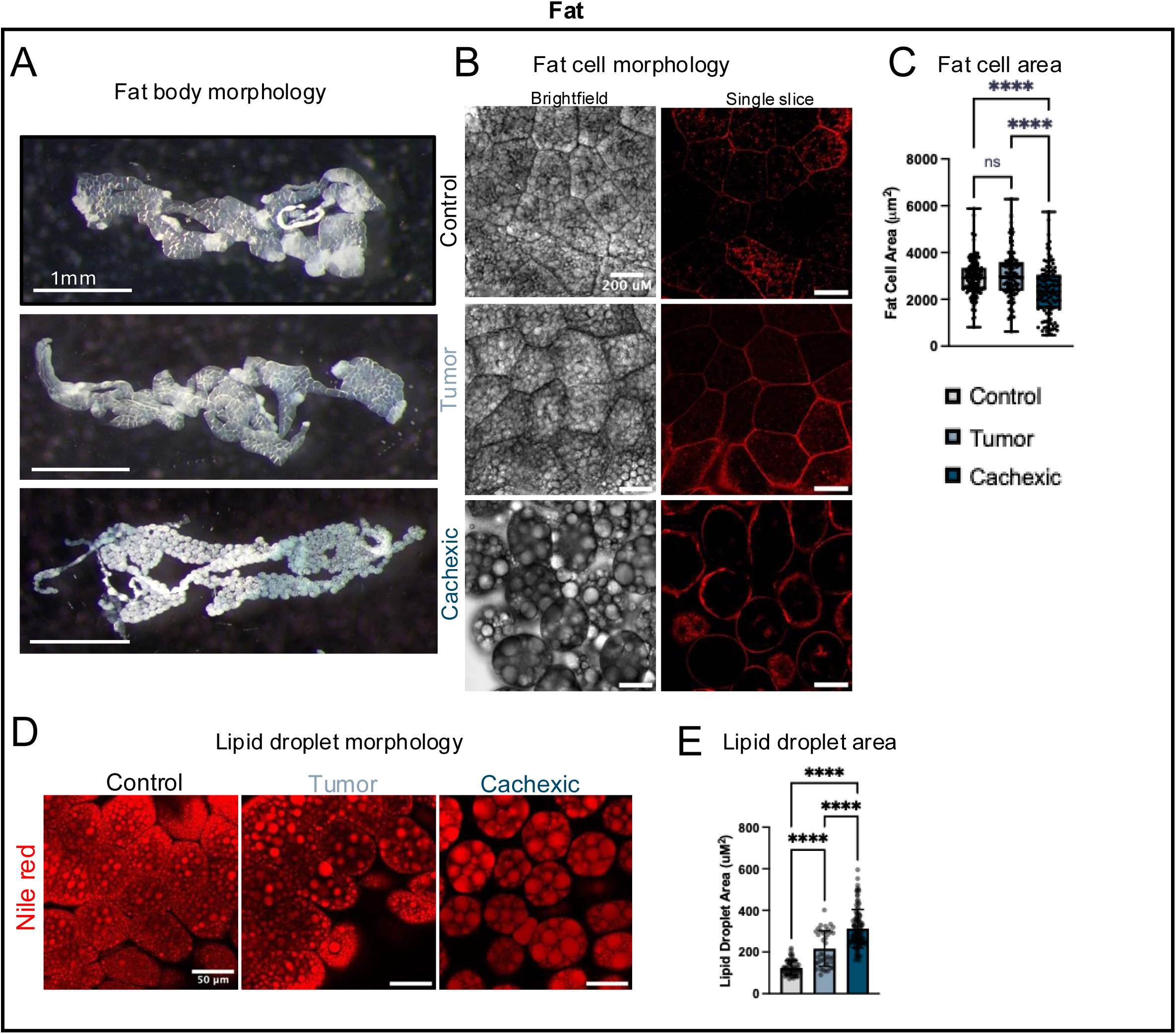
*dpp>Hipk+Sik3-CA* induces morphological and functional changes in the fat body. (A) Dissections of fat body from indicated genotypes of third-instar larvae (n=10/genotype). (B) Single slice brightfield and confocal images of F-actin staining (red). (n=15, 24 and 22) (C) Quantification of fat cell size (n=147, 118 and 116) ns=0.5963, ^∗∗∗∗^p < 0.0001 (Brown-Forsythe and Welch ANOVA test). (D) Representative confocal images of fat body stained with Nile red (n=24, 10 and 29). (E) Quantification of the 5 largest lipid droplets within in each immunofluorescence image (n=60, 40, 105) ∗∗∗∗p < 0.0001 (Brown-Forsythe and Welch ANOVA test).

Drosophila store body fat reserves in the form of lipid droplets in the fat body cells. Due to changes in fat morphology, we hypothesize that there would also be changes in lipid droplet morphology. We found that cachexic larvae had significant lipid droplet accumulation compared to control and tumor-bearing (Fig. 2D-E).

### Functional changes to cachexic larval metabolism results in increased lipolysis and trehalose

The Drosophila fat body is the main site of metabolism, similar to the human liver (Meschi & Delanoue, 2021). Metabolism is the process of using nutrients to produce adenosine triphosphate (ATP) and analogous to humans, flies use carbohydrates, lipids and proteins as energy sources (Chatterjee & Perrimon, 2021). Dietary fatty acids are taken up by the fat body and synthesized and stored as triacylglycerol (TAG), while hemolymph glucose is taken up and condensed into glycogen (for storage) or trehalose, and trehalose is subsequently released into the hemolymph (Chatterjee & Perrimon, 2021; Meschi & Delanoue, 2021) (Fig. 3A). Glucose and trehalose (a non-reducing disaccharide) are the main circulating carbohydrates in the Drosophila hemolymph, with trehalose concentrations more than a 100-fold of glucose (Ugrankar et al., 2015; Wyatt, 1961). Additionally, trehalose has been found to act as a buffer for glucose homeostasis (Matsushita & Nishimura, 2020). As we observed drastic morphological changes in the fat body of cachexic larvae, we wondered if this also resulted in metabolic dysfunction in the cachexic larvae, causing metabolite changes. Whole larval glucose and trehalose concentrations were significantly increased in cachexic larvae (Fig. 3B). This phenotype is found in Drosophila diabetes models and is induced by a high sucrose diet or loss of insulin-like peptides (ILPs) (Rulifson et al., 2002a; Ugrankar et al., 2015). As all the larvae were raised on the same food, yet only the tumor model developed elevated carbohydrate levels, we hypothesize that this phenotype could be mediated by ILPs.

**Figure 3.**
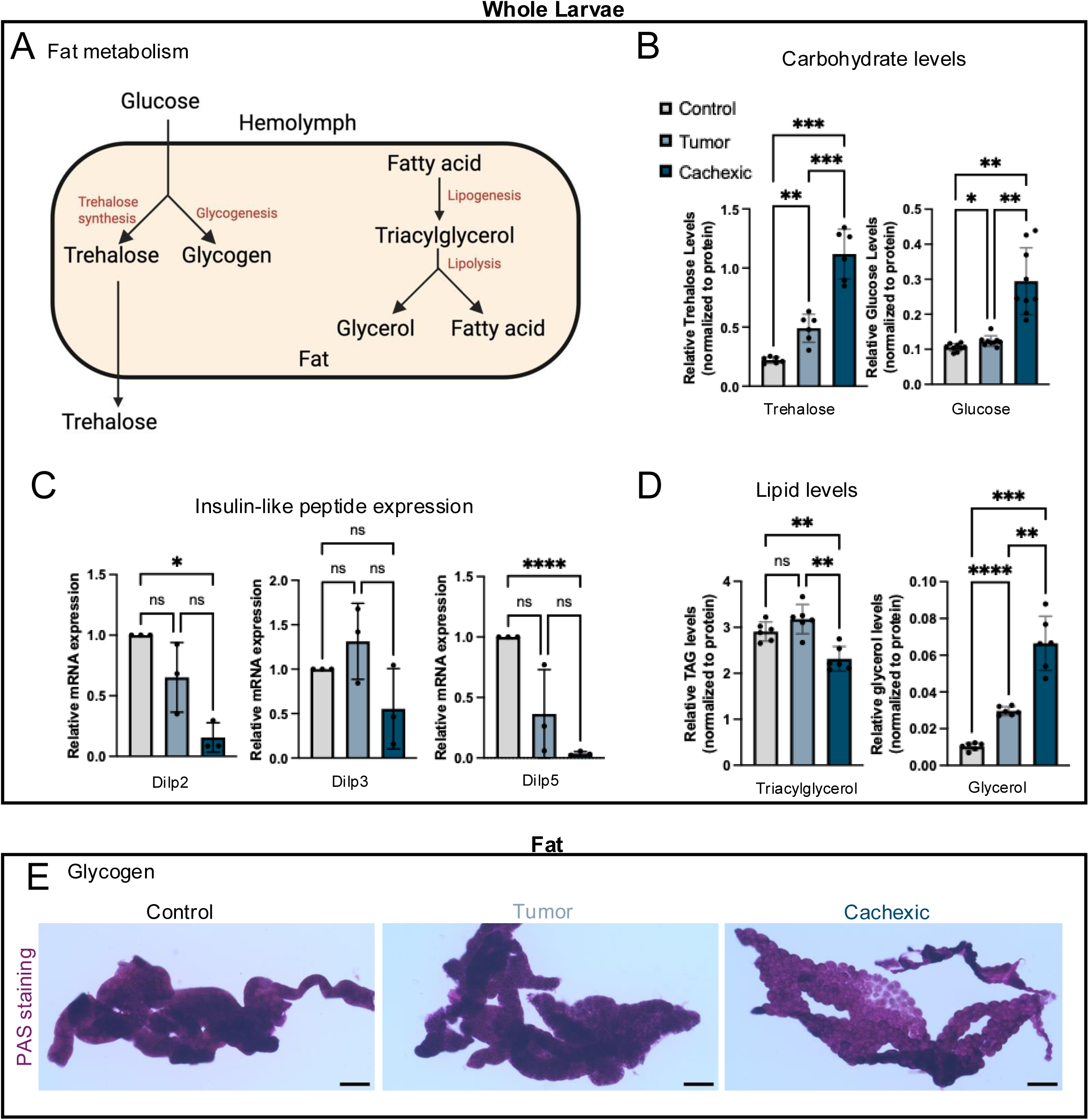
Functional changes to cachexic larval fat results in increased lipolysis and trehalose. (A) Schematic diagram of fat metabolism showing the hemolymph surrounding the fat body (B) Trehalose levels in whole larvae normalized to protein levels (n=5 larvae/data point) ^∗∗^p = 0.0078 and ^∗∗∗^p = 0.0004 and 0.0006 respectively (Brown-Forsythe and Welch ANOVA test). Glucose levels in whole larvae normalized to protein levels (n=5 larvae/data point) ^∗^p = 0.0291 and ^∗∗^p = 0.001 and 0.002 respectively (Brown-Forsythe and Welch ANOVA test). (C) Whole larval *Dilp2, 3* and *5* mRNA levels measured by qPCR (n=5 larvae/data point) ns=0.2938, 0.0895, 0.5315, 0.3847, 0.1601, 0.1683 and 0.4095 respectively, ^∗^p = 0.0125 and ^∗∗∗∗^p < 0.0001 (RM one-way ANOVA). (D) Triacylglycerol levels in whole larvae normalized to protein levels (n=5 larvae/data point) ns=0.2927, ^∗∗^p = 0.0058, 0.0014 respectively (Brown-Forsythe and Welch ANOVA test). Glycerol levels in whole larvae normalized to protein levels (n=5 larvae/data point) ^∗∗^p = 0.0052, ^∗∗∗^p = 0.0007, ^∗∗∗∗^p < 0.0001 (Brown-Forsythe and Welch ANOVA test). (E) Stored glycogen in each tissue was visualized by PAS staining (n=11, 11 and 13 for fat body) scale bar=200uM.

Drosophila ILPs have sequence similarities to human insulins and Drosophila ILP5 (dILP 5) has been shown to activate and bind to the human insulin receptor (Sajid et al., 2011, p. 5). Of the dILPs 1-8, dILPs 2, 3 and 5 are produced and secreted by the insulin producing cells (IPC) in the brain and regulate trehalose levels in Drosophila larvae (Nässel et al., 2015). We find that the mRNA levels of *dILP2* and *dILP5* are significantly reduced in cachexic larvae compared to control (Fig. 3C). Interestingly, *dILP2*, *3* and *5* mutants have increased lipid and glycogen stores (Grönke et al., 2010), while we observed potential fat body wasting which suggests a depletion of stores. Consistent with the fat body wasting, whole larval TAG is significantly reduced and glycerol is significantly increased in cachexic larvae compared to both control and tumor-bearing (Fig. 3D), indicating that even though dILPs 2 and 5 are transcriptionally repressed, lipolysis is increased through a different pathway(s).

As we observed a reduction in TAG levels in cachexic larvae, and we know that lipids exist in the form of intramyocellular lipid droplets (IMLDs) in muscle and are a source of ATP generated from oxidative phosphorylation in the muscle mitochondria. Taken together with reduction in mitochondria density in cachexic larval muscle, we investigated muscle lipid metabolism by staining for lipid droplets in the larval body wall muscle and found approximately a 100 fold reduction in number of lipid droplets in tumor and cachexic larval muscle (Fig. S3A-B), suggesting a reduction of lipid as a fuel source. Interestingly, even though we observed a significant reduction in IMLD number, there was a significant increase in the size of the lipid droplets (Fig. S3C), this could be due to impaired oxidative phosphorylation in tumor and cachexic larval muscle mitochondria, because we observed reduced muscle mitochondria.

In Drosophila larvae, glycogen is mostly stored in the brain, fat body and muscles (Yamada et al., 2018). Under starvation conditions, fat body glycogen is almost completely depleted from the fat body within 4 hours to maintain circulating carbohydrate levels in the larvae, while no change is observed in the brain or muscle glycogen stores (Yamada et al., 2018). We find a slight depletion of glycogen from the fat body of cachexic larvae compared to control and tumor-bearing larvae (Fig. 3E), suggesting that some glycogen might be broken down into glucose as the tumor progresses, potentially contributing to the high trehalose levels.

Altogether, these findings reveal that tumors in epithelial discs induced by overexpression of Hipk and Sik3-CA causes distant organ wasting phenotypes in muscle and fat body.

### Reduction in Lsd2 allows bmm-GFP more access to lipid cachexic larval lipid droplets contributing to increased lipolysis

Drosophila has a dual lipolytic system: the Adipokinetic hormone (Akh) signaling pathway and the Brummer lipase (Grönke et al., 2007). Because we observe increased lipolysis in cachexic larvae, we hypothesized that one or both lipolytic systems are activated to mediate this observed depletion of TAG levels. Drosophila have only two perilipins, PLIN1/Lsd-1 and PLIN2/Lsd-2 and they have both been found to protect the lipid droplets from lipolysis mediated by Brummer (Bmm) lipase, the Drosophila homolog of human Adipose triglyceride lipase (ATGL) (Bi et al., 2012; Grönke et al., 2005). Therefore, we tested the levels of perilipins in cachexic larvae. Lsd1 protein levels is not significantly changed in the fat body, while Lsd2 protein levels are significantly reduced in cachexic larvae compared to tumor-bearing (Fig. 4A-B). To determine if reduction in Lsd2 protein levels is correlated with reduced Lsd2 protein association with lipid droplets, we examined Lsd2 localization in fat cells. In both control and tumor fat cells, Lsd2 preferentially associates with smaller peripheral lipid droplets as indicated by the higher Lsd2 fluorescence gray value peaks (Fig. 4C), consistent with previous research (Bi et al., 2012). However, in cachexic larvae, Lsd2 seems to have less association with peripheral lipid droplets and is more diffuse throughout the fat cell cytoplasm (Fig. 4C). As Lsd2 protects lipid droplets from Brummer-mediated lipolysis, we examined the cellular localization of Bmm lipase with an endogenous C-terminal GFP knockin (Zhao et al., 2022). In control larval fat, Bmm-GFP accumulates around mostly bigger lipid droplets, with low levels in the cytoplasm of fat cells (Fig. 4D). In both tumor and cachexic larval fat, Bmm-GFP is more accumulated around both bigger and smaller peripheral lipid droplets and there also seems to be more cytoplasmic Bmm-GFP (Fig. 4D). Quantification of Bmm-GFP fluorescence intensity in fat cells is consistent with this observation and reveals that tumor and cachexic larvae has significantly more average Bmm-GFP fluorescence intensity in fat cells than control (Fig. 4E). Quantification of larval fat Bmm-GFP corroborates this observation, tumor and cachexic larval fat Bmm-GFP protein levels are significantly increased compared to control (Fig. 4F-G).

**Figure 4.**
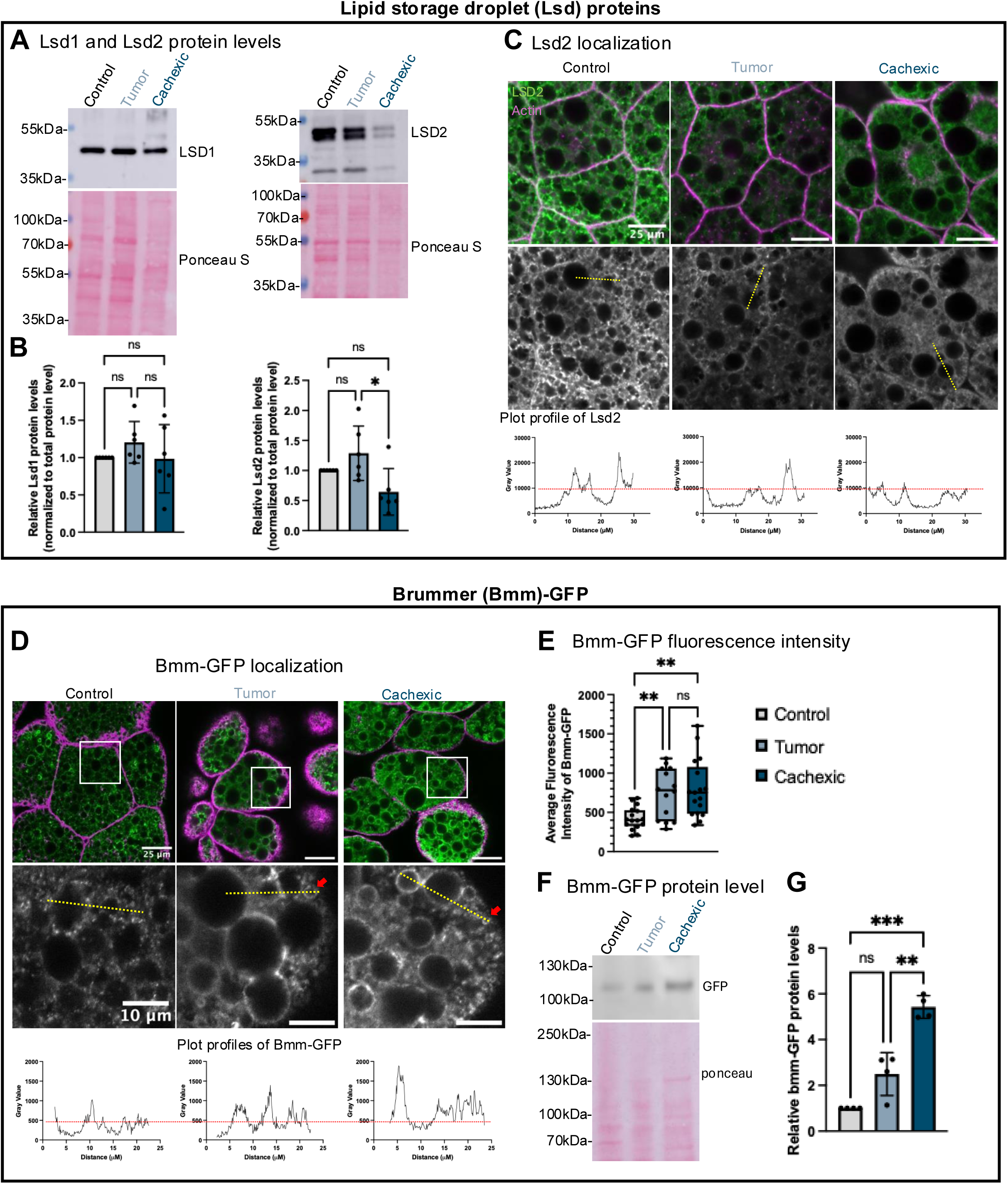
Reduction in Lsd2 allows bmm-GFP more access to cachexic larval lipid droplets. (A) Western blot of whole protein extracts of fat tissue from third instar larvae probed for the presence of Lsd1 and Lsd2 protein. Ponceau S is used as a loading control. n=6 biologically independent experiments. (B) Lsd1 and Lsd2 protein levels were quantified and normalized to control. Ponceau S-stained membranes were used to verify total protein loading in each sample. n=10 larval fat bodies/data point ns=0.2635, 0.9964, 0.3807, 0.3455, 0.156 and ^∗^p = 0.0426 (ANOVA Tukey’s multiple comparisons test). (C) Representative single slice confocal images of F-actin staining (fuchsia) and LSD2 (green). Greyscale pictures of Lsd2 channel are shown below. Line graphs of plot profiles (yellow dotted line) showing distribution of fluorescence intensity of LSD2 from respective tissues shown above (n=18, 14 and 16). (D) Representative single slice confocal images of F-actin staining (fuchsia) and Bmm-GFP (green). Magnified sections of Bmm-GFP channel (grey) is shown below. Higher Bmm-GFP protein accumulation around smaller lipid droplets is indicated by the red arrows. Line graphs of plot profiles (yellow dotted line) showing distribution of fluorescence intensity of Bmm-GFP from respective tissues shown above. (E) Quantification of average fluorescence intensity of fat Bmm-GFP levels. (n=16, 14 and 17) ns=0.91697, ^∗∗^p = 0.0071 and 0.0027 respectively (Brown-Forsythe and Welch ANOVA test) (F) Western blot of whole protein extracts of fat tissue from third instar larvae and probing for the presence of Bmm-GFP protein. Ponceau S is used as a loading control. *n* = 4 biologically independent experiments (G) Bmm-GFP protein levels were quantified and normalized to control. Ponceau S-stained membranes were used to verify total protein loading in each sample. n=10 larval fat bodies/data point ns=0.0979, **p=0.0053 and ***p = 0.0008 (ANOVA Tukey’s multiple comparisons test).

### Glucagon-like Adipokinetic hormone (Akh) production is upregulated in *dpp>Hipk+Sik3-CA* tumor-bearing larvae

*Akh* is exclusively expressed in the corpora cardiaca (CC) cells of the ring gland (RG) in Drosophila larvae (Kim & Rulifson, 2004; Lee & Park, 2004) (Fig. 5A). Additionally, ectopic expression of *Akh* induces increased lipolysis and trehalose levels in Drosophila larvae (Lee & Park, 2004), similar to our observations in cachexic larvae. Therefore, we examined transcription levels of *Akh* in the brain and RG tissue and found that *Akh* is significantly upregulated in tumor-bearing larvae (Fig. 5A). We also observed increased Akh fluorescence intensity and CC cell area in both tumor-bearing and cachexic larvae (Fig. 5B-D), indicating that there are increased Akh peptide levels in these larvae. Taken together, this indicates that both Bmm and Akh lipolytic pathways are upregulated to induce increased trehalose and lipolysis in cachexic larvae.

**Figure 5.**
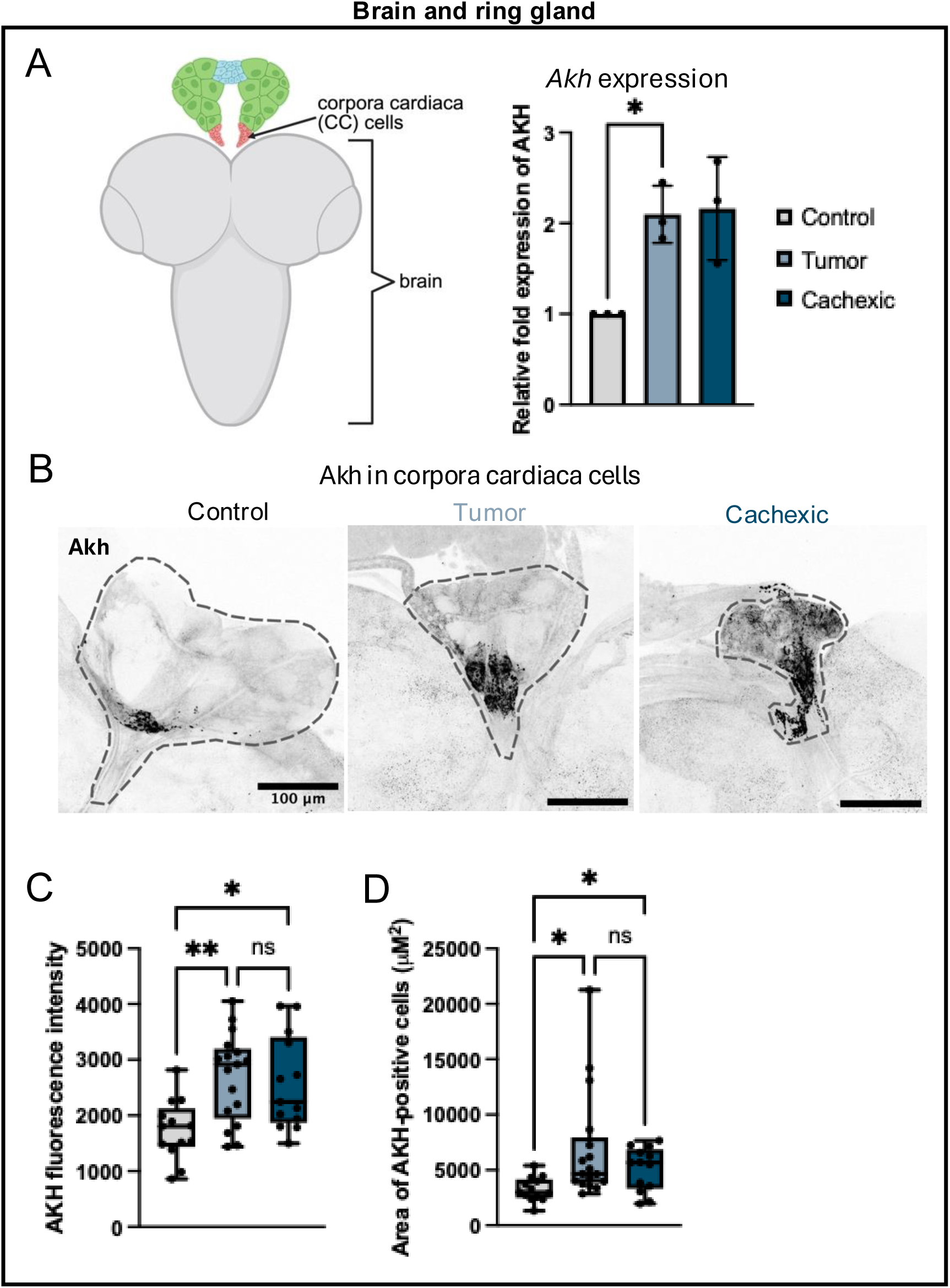
Glucagon-like Adipokinetic hormone (Akh) production is upregulated in *dpp>Hipk+Sik3-CA* tumor-bearing larvae. (A) Schematic diagram of larval brain and ring gland (left) and brain and prothoracic gland AKH expression level (right) (*n*=3, 5 flies/biological replicate), *p=0.0265, ns=0.0714 (Welch’s t-test) (B) Maximal projection images of the ring gland (circled in the dotted line) of female L3 larvae immunostained for AKH (black, inverted LUT). (C) Quantification of AKH fluorescence intensity (n = 13, 13 and 17 respectively). *n*=3 ns=0.9794, ^∗^p = 0.02 and ^∗∗^p = 0.0017 (Brown-Forsythe and Welch ANOVA test). (D) Quantification of AKH signal area (uM^2^) (n = 13, 13 and 17 respectively). *n*=3 ns=0.4647 and ^∗^p = 0.0268 and 0.0238 respectively (Brown-Forsythe and Welch ANOVA test).

Therefore, we wondered if a reduction in the Akh lipolytic pathway would reduce the aberrant metabolism in tumor and cachexic larvae and subsequently fat wasting. Hence, we induced tumors in either a heterozygous *Akh^AP^* or *Akh receptor (AkhR^1^)* mutant background. Tumor and cachexic larvae heterozygous for *Akh^AP^* did not show any effects on tumor proliferation (Fig. S4A). Heterozygosity for *Akh^AP^* did significantly reduce trehalose levels in tumor larvae but did not rescue the increased trehalose and lipolysis in cachexic larvae (Fig. S4B-C). Interestingly, *AkhR^1^* heterozygous cachexic larvae are significantly smaller in terms of protein levels than cachexic larvae (*dpp> Hipk +Sik3-CA* D16 larvae) (Fig. S4D-E), even though they have the same aberrant metabolism, specifically increased trehalose and lipolysis (Fig. S4F-G), indicating that cachexic larval size is a separate paraneoplasia from organ wasting.

### Tumor-derived Unpaired ligands induce Glucagon-like Adipokinetic hormone (Akh) production in *dpp>Hipk+Sik3-CA* tumor-bearing larvae through upregulation of the Jak/Stat signaling pathway

Tumors use external sources of nitrogen and carbon to sustain uncontrolled cell proliferation, and therefore metabolic reprogramming of the host organism by tumor secreted factors to promote tissue wasting supports cancer growth (Porporato, 2016). Consequently, the tumors overexpressing Hipk and Sik-CA in our model could potentially be secreting cachexic factors to induce Akh and Bmm-GFP, resulting in increased lipolysis and the production of trehalose. A previous study in adult Drosophila found that the Unpaired 2 (Upd2) cytokine controls Akh secretion (Zhao & Karpac, 2017). To determine if Unpaired ligands are upregulated in our tumor model, we quantified the transcription levels of unpaired ligands 1, 2 and 3 and found that they are all significantly upregulated approximately 5-15 times in both tumor and cachexic larval tumors (Fig. 6A).

**Figure 6.**
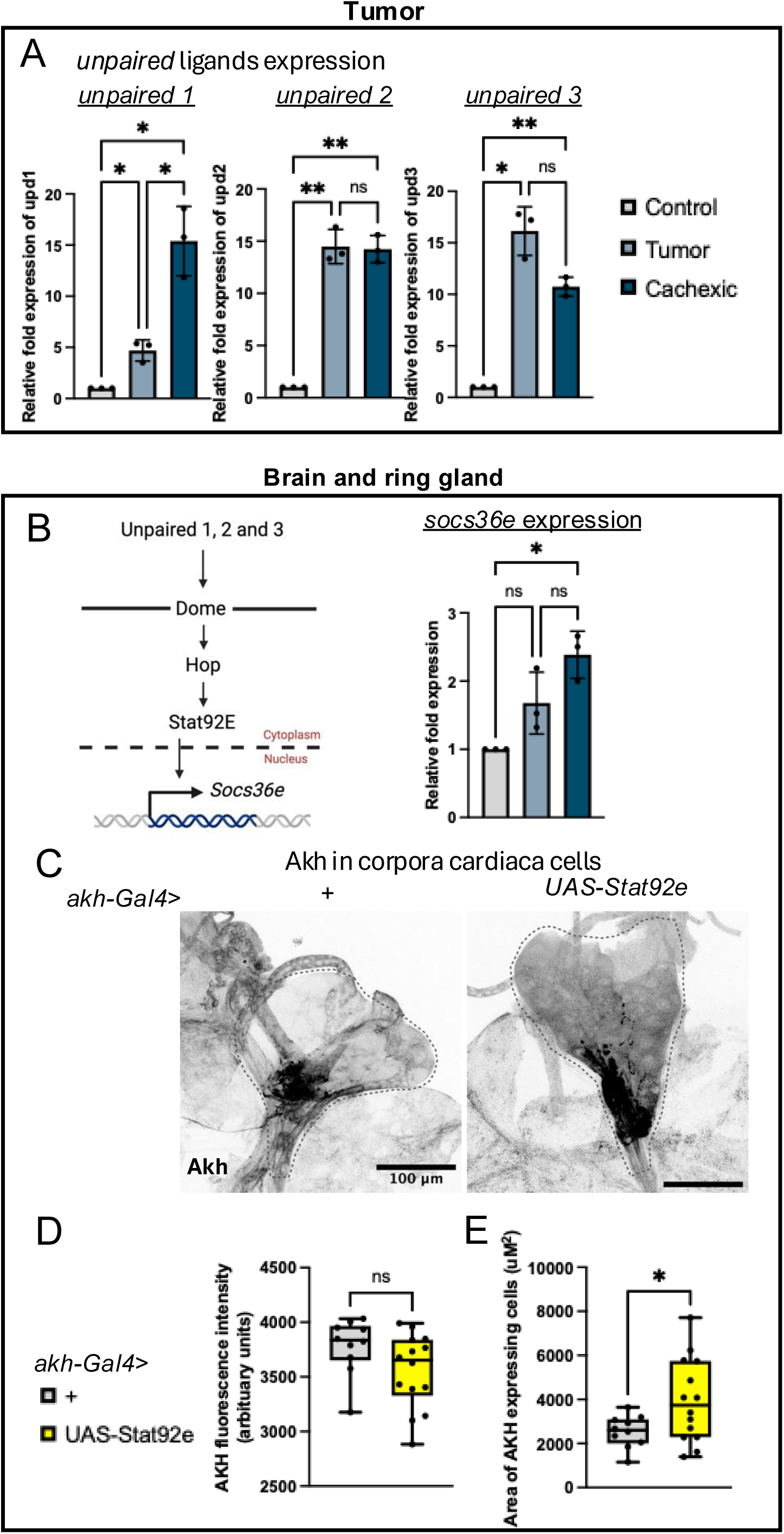
Tumor-derived unpaired ligands induce Glucagon-like Adipokinetic hormone (AKH) production in *dpp>Hipk+Sik3-CA* tumor-bearing larvae through upregulation of the Jak/Stat signaling pathway. (A) Schematic diagram of Jak/Stat signaling pathway (left) and epithelial discs *unpaired 1, 2 and 3* expression level (right) (n=3, 5 flies/biological replicate), *p=0.0455, 0.0327, 0.0309 and 0.0145, **p=0.0089, 0.0058 and 0.0052, ns=0.9858 and 0.1774 respectively (B) Brain and prothoracic gland *socs36e* expression level (*n*=3, 5 flies/biological replicate), *p=0.0363, ns=0.2139 and 0.1689 respectively (RM one-way ANOVA). (C) Maximal projection images of the ring gland (circled in the dotted line) of female L3 larvae immunostained for AKH (black, inverted LUT). (D) Quantification of AKH fluorescence intensity (n = 10 and 14 respectively). ns=0.0783 (Welch’s t test). (E) Quantification of AKH signal area (uM^2^) (n = 10 and 14 respectively). ^∗^p = 0.0215 (Welch’s t test).

Upd ligands activate the Jak/Stat pathway by binding to Dome receptors which activate the Jak Hopscotch (Hop). Activated Hop phosphorylates Stat92E which then dimerizes and translocates to the nucleus to drive target gene expression (Harrison et al., 1998; Hou & Perrimon, 1997) (Fig. 6B). To determine if these Unpaired ligands are secreted and activating the Jak/Stat pathway in CC cells, we used *socs36e* as a transcriptional readout of the Jak/Stat pathway and found that it was significantly induced in the brain and ring gland of cachexic larvae (Fig. 6B), indicating that the upregulation of Akh in tumor and cachexic larvae could be a result of activation of Jak/Stat signaling in cc cells. Finally, overexpression of the Jak/Stat transcription factor Stat92E using a CC specific *akh-Gal4* driver resulted in increased Akh signal area (Fig. 6C-E), consistent with results found in adult Drosophila (Zhao & Karpac, 2017). Taken together, these results suggests that the tumor is potentially secreting unpaired ligands into the hemolymph, and activating Jak/Stat signalling pathway in the cc cells, inducing increased Akh peptide levels.

## Discussion

In this study we establish that overexpression of Hipk and Sik3-CA in epithelial tissues of Drosophila larvae produce a novel Drosophila cancer cachexia model. We also establish tumour-peripheral organ communication that results in changes in host metabolism. In particular, our results identify an Upd-Akh signaling axis that contributes to increased trehalose levels and lipolysis in the cachexic larvae (Fig. 7). As the Drosophila Jak/Stat and glucagon-like Akh signaling pathways are highly conserved in humans, we believe that this novel Drosophila cancer cachexia model could provide more insight into the complex metabolic changes in human cancer cachexia.

**Figure 7.**
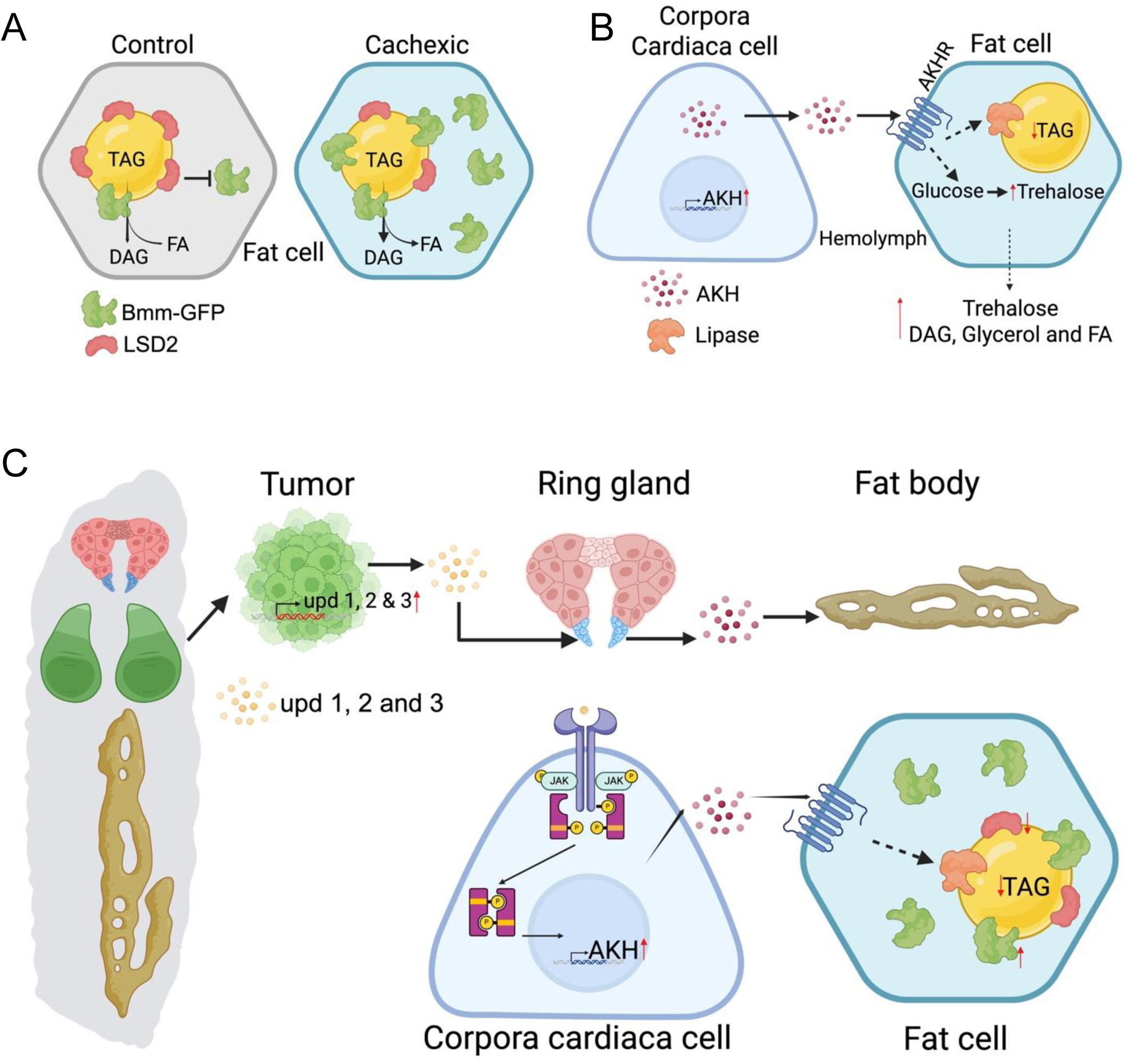
Model of changes in distant organ functions and metabolite changes induced by overexpression of Hipk and Sik3-CA in larval epithelial discs. (A-C) This graphical abstract was created using Biorender.

We started this line of investigation due to the developmental delay phenotype that we saw in our cancer cachexia model, that is also commonly seen in other Drosophila cancer and cancer cachexia models (Liu et al., 2022; Menut et al., 2007). It has been established that this developmental delay is caused by an upregulation of insulin-like peptide 8 (DILP8) and Unpaired ligands in response to injury or tumor, which inhibits ecdysone synthesis hence delaying metamorphosis (Cao et al., 2022; Colombani et al., 2012; Garelli et al., 2012). From various studies, developmental delay by itself is not sufficient to induce distant organ wasting (Hodgson et al., 2021), and reduction or silencing *DILP8* in Drosophila cancer cachexia models rescues developmental delay without rescuing cancer cachexia (Santabárbara-Ruiz & Léopold, 2021; Yeom et al., 2021). Therefore, developmental delay and cancer cachexia are separate paraneoplasias.

We found muscle dysfunction and wasting in our cancer cachexia model, consistent with the *scrib Ras^V12^* and *yki^act^* Drosophila cancer cachexia models (Figueroa-Clarevega & Bilder, 2015; Kwon et al., 2015). This is commonly found in human cancer cachexia patients as well, and it is usually caused by multiple interconnected signaling pathways that converge to induce muscle wasting through activation of catabolic pathways (Rausch et al., 2021; Setiawan et al., 2023). Of relevance, mitochondrial dysfunction is commonly associated with cancer cachexia (Setiawan et al., 2023). Interestingly, excessive fatty acid oxidation in mouse skeletal muscle was found to induce muscle wasting under a lipid-induced insulin resistance disease condition (Koves et al., 2008). Additionally, in the *Ras^V12^dlg^RNAi^* Drosophila cancer cachexia model, muscle wasting was mediated by increased lipid utilization due to mitochondrial fusion (Dark et al., 2024). This could explain why we found reduced number of lipid droplets in the tumor and cachexic larval muscle, due to increased breakdown of lipids into fatty acids and subsequent fatty acid oxidation. It would be interesting to test whether in our model the larval muscle is also insulin resistant. In contrast to our finding, studies in human, mouse and Drosophila muscle have found increased number and size of lipid droplets in cancer cachexia (Dark et al., 2024; Hodgson et al., 2021; Stephens et al., 2011). This increase in lipid droplets is hypothesized to be due to reduced fat oxidation in muscle mitochondria. Either way, both these phenotypes point to mitochondrial dysfunction in cachexic muscle. In addition, interleukin-6 (IL-6) the mammalian homolog of unpaired 3, is a known myokine that induces muscle wasting (Agca & Kir, 2024; Muñoz-Cánoves et al., 2013). This is also consistent with the *yki^act^* Drosophila cancer cachexia model in which the authors showed that gut tumors secrete Upd3 to induce muscle wasting (Ding et al., 2021).

We find that the cachexic larval fat body to be the central regulator of metabolite changes in this Drosophila cancer cachexia model. Morphologically, we also find increased lipid droplet size associated with tumor and cachexic larvae, consistent with findings from the *scrib Ras^V12^* and *Ras^V12^dlg^RNAi^* Drosophila cancer cachexia models (Figueroa-Clarevega & Bilder, 2015; Lodge et al., 2021). Of note, reduction in PLIN2/Lsd2 and increased ATGL/Bmm results in smaller lipid droplets due to increased lipolysis (Beller et al., 2010; Grönke et al., 2005). Lipid droplet size is determined by the lipid content, composition of the phospholipid monolayer, and lipid-droplet associated proteins or perilipins (Zhao et al., 2022). This suggests that the larger lipid droplets we observe in our cachexia model are the result of changes to lipid content or phospholipid composition.

Metabolically, the functional changes such as increased carbohydrates and lipolysis are consistent with other Drosophila, mice and human cancer cachexia models (Agustsson et al., 2007; Berriel Diaz et al., 2024; Dev et al., 2018). In human cancer cachexia patients, the increase in glucose is proposed to be a result of lactate secreted from tumor cells leading to increased hepatic glucose production (Masi & Patel, 2021). Additionally, adipose tissue loss in human patients is a result of increased lipolysis and lipid utilisation, and potentially impaired lipogenesis (Fang et al., 2023).

Of note, insulin resistance is a common co-morbidity in cancer cachexia models and patients (Dev et al., 2018; Figueroa-Clarevega & Bilder, 2015; Khezri et al., 2021; Kwon et al., 2015). However, in our model, we found that hyperglycemia is caused by a reduction in dILPs or a suppression of the insulin pathway. It should be noted that in a study with over 600 cancer patients, over a third showed insulin resistance (Glicksman & Rawson, 1956), indicating that around two thirds of patients did not have insulin resistance, pointing to the heterogeneity of cancer effects. This highlights the necessity of characterizing different models of cancer cachexia as human cancer cachexia is a complicated and heterogenous disease.

We find that elevated Bmm-GFP and Akh mediate increased lipolysis in cachexic larvae (Fig. 7A-B). This is consistent with a mouse model of cancer cachexia in which they found *Adipose triglyceride lipase* (*Atgl)*-deficient mice (ATGL is the mammalian homolog of Bmm) with tumors rescued the increased lipolysis, and adipose tissue and muscle wasting (Das et al., 2011). To further address mechanisms underlying the observed phenotypes, future experiments could incorporate an additional expression system along with UAS-Gal4 to induce a fat specific knockdown of *brummer* in cachexic larvae to investigate any reduction in lipolysis and rescue of distant organ wasting phenotypes. As we also found increased Akh in tumor and cachexic larvae, there are also numerous mouse and human cancer cachexia studies showing increased glucagon (mammalian functional homolog of Akh) plasma levels (Bartlett et al., 1995; Ding et al., 2025). Our model shows that Unpaired ligands are upregulating the Jak/Stat signaling pathway in CC cells and subsequently upregulating Akh to induce lipolysis (Fig. 7C). Recently, another study also showed that Akh is essential for host distant organ wasting in the yki^3SA^ gut tumor model, consistent with this paper’s findings (Ding et al., 2025). However, they found that PDGF- and VEGF-related factor (Pvf1) to be the cachexic factor mediating Akh upregulation (Ding et al., 2025). Given that the tumor has been shown to secrete numerous cachexic factors to promote distant organ wasting (Liu et al., 2022; Setiawan et al., 2023), multiple factors could be contributing to the same phenotype. Furthermore, Upd3/IL-6 is often associated with cancer cachexia in both Drosophila, mouse and human cancer cachexia (Bonetto et al., 2012; Ding et al., 2021). While these studies have mostly focused on muscle wasting induced by Upd3/IL-6 inflammation, our study shows a novel connection between Upd ligands activating lipolysis through Akh. Outside of the context of cancer cachexia, this result is consistent with human and mouse research, which found that IL-6 stimulates glucagon secretion (Chow et al., 2014). It would be interesting to know if in the context of cancer cachexia, IL-6 can also stimulate glucagon secretion in human patients.

While trying to reduce Akh signaling, and therefore lipolysis, by inducing the Hipk + Sik tumor in heterozygous *Akhr* mutants, we observed that the cachexic larvae in the heterozygous *Akhr* mutant background are smaller in size and therefore had significantly less protein levels compared to cachexic larvae. However, the reduction in size had no effect on the increased lipolysis and trehalose levels, indicating that increased larval size in this context is a separable non-autonomous effect from organ wasting. Regulation of larval body or organ size, is dependent on the environment, such as nutrition and temperature, and inter-organ coordination mostly through hormones and secreted proteins (Mirth & Shingleton, 2012). Adult *Akhr* mutants do not have any significant changes in size (Bharucha et al., 2008), therefore the reduction in *Akhr* in cachexic larvae is not likely to contribute to smaller larvae. Ablation of IPCs which secrete the dILPs 2, 3 and 5 reduced larvae size (Rulifson et al., 2002b), which is inconsistent with our cancer cachexia model as we see reduced *dILP 2* and *5* expression levels, suggesting that the reduction in cachexic larval size is due to other factors.

Our study shows that tumors are secreting Unpaired ligands to upregulate Akh in the CC cells, resulting in increased trehalose and lipolysis in the cachexic larvae. Tumors use external sources of nitrogen and carbon to sustain uncontrolled cell proliferation, and therefore metabolic reprogramming of the host organism by tumor secreted factors to promote tissue wasting supports cancer growth (Porporato, 2016). Therefore, we speculate that this organismal change in metabolites (increase in carbohydrates and fatty acids) potentially feeds into tumor growth. Thus, we are currently investigating the energy metabolism of tumors, such as rates of glycolysis and oxidative phosphorylation. Of note, a previous study suggests that Sik3 acts cell-autonomously downstream of both the insulin and Akh pathways to regulate *bmm* expression in the fat body (Choi et al., 2015). Please note that in our study Sik3-CA is overexpressed in the epithelial discs and not in the fat body, therefore Sik3-CA is regulating insulin and Akh pathway potentially with Hipk non-autonomously. It would be of interest to test whether *Sik3-CA* overexpression in the tumors is increasing *bmm* expression and if so, if the tumor is utilizing the fatty acid released from lipolysis for increased proliferation. In addition, cancer cells’ increased glutamine consumption is correlated with Myc activation in human cancer cell lines and mouse cancer models (Zheng, 2012). As we also found increased dMyc protein levels in the Hipk + Sik tumors (Yu et al., 2023), we could test if there is also increased glutamine consumption contributing to the increased cell proliferation mediated by increased dMyc.

## Material and methods

### Drosophila culture

*Drosophila melanogaster* flies were raised on standard cornmeal-molasses food. Stocks were kept at 18 °C or room temperature. Crosses were carried out at 29 °C as indicated. The following fly strains were used: *dpp-Gal4/TM6B* (Staehling-Hampton et al., 1994), *akh-gal4* (BL #25684), *UAS-GFP* (BL #5431), *UAS-RFP* (BL #7119), *UAS-Hipk^3M^* (Blaquiere et al., 2018), *UAS-Sik3-CA* (Wehr et al., 2013), Bmm-GFP (Zhao et al., 2022), *w^1118^* (BL #5905), *UAS-Stat92E-GFP* (Sotillos et al., 2013), *Akh^AP^* (Gáliková et al., 2015), and *AkhR1* (Grönke et al., 2007). Strains with the BL stock number were obtained from the Bloomington Drosophila Stock Center (Bloomington, IN, USA).

### Larval mass and hemolymph volume

We adapted the protocol from (Santabárbara-Ruiz & Léopold, 2021). Third instar larvae were first rinsed in PBS 1X and dried on a Kimwipe. They were then weighed in groups of 5 to determine mass. Groups of 5 larvae were transferred to parafilm, and abdominal openings were made with forceps, allowing the hemolymph to pool out surrounding the larvae. Hemolymph was then extracted with a pipette tip.

### Pupal counting

Flies were allowed to lay eggs for 24 hours. We then counted number of pupae on the walls of the food vials of our genotypes of interest that pupated over 14 days.

### Immunohistochemistry

Imaginal discs from late L3 larvae were dissected in phosphate-buffered saline (PBS) and fixed in 4% paraformaldehyde (PFA) for 15 min at room temperature. After fixation, samples were washed in PBS with 0.1% Triton X-100 (PBST). After blocking with 5% BSA (Bovine serum albumin) in PBST for 1 h at room temperature, samples were incubated with primary antibodies overnight at 4°C.

Larval body wall muscle dissection was adapted from (Dark et al., 2022). Third instar larvae are washed in PBS and subsequently heat killed in 70°C–75°C PBS for 5 seconds. Larval muscle fillets are dissected and fixed in ice cold 4% PFA for 20 minutes at room temperature. To stain for lipid droplets in muscle, larval body wall muscle was incubated overnight at 4 °C in Bodipy 493/503 (Invitrogen D3922).

Larval fat dissections was adapted from (Dark et al., 2022). Third instar larvae are dissected in PBS and fixed in 4% PFA for 30 minutes at room temperature. To stain for lipid droplets, fixed fat tissue were incubated for 30 minutes at room temperature in Nile Red dye (1:500) (Invitrogen #N1142). After mounting, fat samples are imaged immediately or within the same day.

The primary antibodies used include rabbit anti-Hipk (1:200) (Blaquiere et al., 2018), mouse anti-atp5a (1:500) (Abcam ab14748), rat anti-Dcad2 (1:100) (DSHB AB_528120), mouse anti-MMP1 (1:100) (DSHB 3A6B4, 3B8D12 and 5H7B11 used together), rabbit anti-LSD2 (1:500) (gift from Dr. Ronald Kühnlein), rabbit anti-GFP (1:500) (Thermo Fisher A11122), rabbit anti-AKH (1:500) (gift from Dr. Wei Song).

After washing with PBST (PBS for Nile red staining and PBS in 0.05% Saponin for Bodipy staining of muscle), samples were incubated with FITC and/or Alexa Fluor 647-conjugated secondary antibodies (1:500, Jackson ImmunoResearch Laboratories, Inc.), DAPI (Invitrogen D1306) was used at 1:500 and Phalloidin-Rhodamine (Invitrogen R415) was used at 1:1000. Samples were mounted in VECTASHIELD® Antifade Mounting Medium (H-1000-10) after wash. Images were taken on a Zeiss LSM880 Airyscan confocal microscope and processed using ImageJ (Schindelin et al., 2012).

### Locomotion Assay

We adapted the protocol from (Fushiki et al., 2016). Third instar larvae were rinsed in PBS 1X and dried on a Kimwipe and then transferred to an agar plate and acclimatized for 2 minutes. The larvae were then videoed using an iPhone 14 Pro. The peristaltic movement was manually measured in the videos over 30 seconds.

### Western blotting

Hemolymph extracted from larvae was diluted 10X in 5X Laemli Buffer. Cells or tissues were lysed with 1× Cell Lysis Buffer (Cell Signaling Technology #9803), supplemented with 1× Protease Inhibitors (Roche), and 1 mM phenylmethylsulfonyl fluoride. All samples were heated at 95°C for 5 minutes.

Lysates were stored at −20°C. Protein lysates were resolved by 4%-15% SDS/PAGE (at 90 V for 120 min) and then transferred to nitrocellulose membranes (at 20 V for 65 min). For Ponceau-S staining, nitrocellulose membranes were incubated in Ponceau-S solution (40% methanol (v/v), 15% acetic acid (v/v), 0.25% Ponceau-S) for 5 minutes at RT. The membranes were destained using TBST. Membranes were blocked with 5% BSA in TBST before primary and secondary antibody incubation. Images were acquired by a FujiFilm LAS-4000 Chemiluminescent Scanner.

The primary antibodies used: mouse anti-Myosin heavy chain (1:500, DSHB Hybridoma Product BB7/86.1), mouse anti-MMP1 (1:500) (DSHB 3A6B4, 3B8D12 and 5H7B11 used together), rabbit anti-LSD1 1:3000 and rabbit anti-LSD2 1:5000 (gift from Dr. Ronald Kühnlein), rabbit anti-GFP (1:5000) (Thermo Fisher A11122).

### Measurement of Carbohydrate and Lipid Levels

5 female larvae were homogenised in 500 uL PBST. The solution was incubated at 75°C for 5 minutes, then spun down at 14,000 rpm for 10 minutes at 4°C. Supernatant was collected and used to perform glucose (FujiFilm Autokit Glucose #997-03001), trehalase (MilliporeSigma #T8778-1UN), TAG (Cayman Triglyceride Colorimetric Assay Kit #10010303-96), glycerol (Cayman Glycerol Colorimetric Assay Kit #10010755-96, Glycerol Colorimetric Assay Kit #10010755-96) and Bradford assays (Thermo Scientific Pierce™ BCA Protein Assay Kit #23225) according to manufacturer’s instructions.

### qRT-PCR

Whole larvae from the third instar larval stage or specific tissue dissected from third instar larvae were briefly washed in PBS and temporarily stored in RNAlater Stabilization solution (Invitrogen AM7020). Total RNA was extracted using RNeasy Mini Kits (Qiagen 74101). The first strand of cDNA was synthesized using the OneScript Plus cDNA Synthesis Kit (Abm G236). qRT-PCR were performed using StepOne Real-time PCR System (Applied Biosystems). Primers used are:

*Dilp2* F: ACGAGGTGCTGAGTATGGTGTGC
*Dilp2* R: CACTTCGCAGCGGTTCCGATATCG

*Dilp3* F: GTCCAGGCCACCATGAAGTTGTGC
*Dilp3* R: CTTTCCAGCAGGGAACGGTCTTCG

*Dilp5* F: TGTTCGCCAAACGAGGCACCTTGG
*Dilp5* R: CACGATTTGCGGCAACAGGAGTCG

*MMP1* F: AGGACTCCAAGGTAGACACAC
*MMP1* R: TTGCCGTTCTTGTAGGTGAACGC

*Akh F*: TCCCAAGAGCGAAGTCCTCA
*Akh R*: CCAGAAAGAGCTGTGCCTGA

*Unpaired 1 F*: TGGATCGACTATCGCAACTTCG
*Unpaired 1 R*: GTCCTGGCTACTGTTTAGGCT

*Unpaired 2 F*: GAGGGCAGCTACGACAGTG
*Unpaired 2 R*: GGAGAAGAGTCGCAGGTTGT

*Unpaired 3 F*: CTGGTCACTGATCTTACTCGCC
*Unpaired 3 R*: GGATTGGTGGGATTGATGGGA

*Socs36e F*: ATGGGTCATCACCTTAGCAAGT
*Socs36e R*: TCCAGGCTGATCGTCTCTACT

### Histochemistry

Larval fat bodies were dissected in PBS and fixed in 4% PFA for 20 minutes at room temperature. Tissues were then washed in PBS and incubated with periodic acid for 5 minutes. To stain for glycogen, fixed fat tissues were incubated for 30 minutes at room temperature in Schiff’s reagent. After mounting, fat samples are imaged immediately or within the same day. The fat tissues were imaged with a Zeiss Axioplan-2 microscope and processed with ImageJ software.

### Statistical Analyses

All statistical analyses were performed using GraphPad Prism 10.3.1 (GraphPad Software, San Diego, USA). Analyzed data with *p* < .05 was considered statistically significant.

## Acknowledgments

We thank Dr. Nicolas Tapon, Dr. Norbert Perrimon and Dr. Ronald Kühnlein for fly strains. We are grateful to Dr. Christopher Beh, Dr. Caterina Ramogida, Dr. Ronald Kühnlein, Dr. Wei Song and the Developmental Studies Hybridoma Bank (DSHB) for providing valuable antibodies and other reagents. This work was supported by operating grants from Natural Sciences and Engineering Research Council of Canada (NSERC, RGPIN/2020-06192) and Canadian Institutes of Health Research (CIHR, PJT-156204). K.Y. was supported by a Simon Fraser University Dean of Graduate Studies Entrance Scholarship.

---

**Supplemental Fig. 1.**
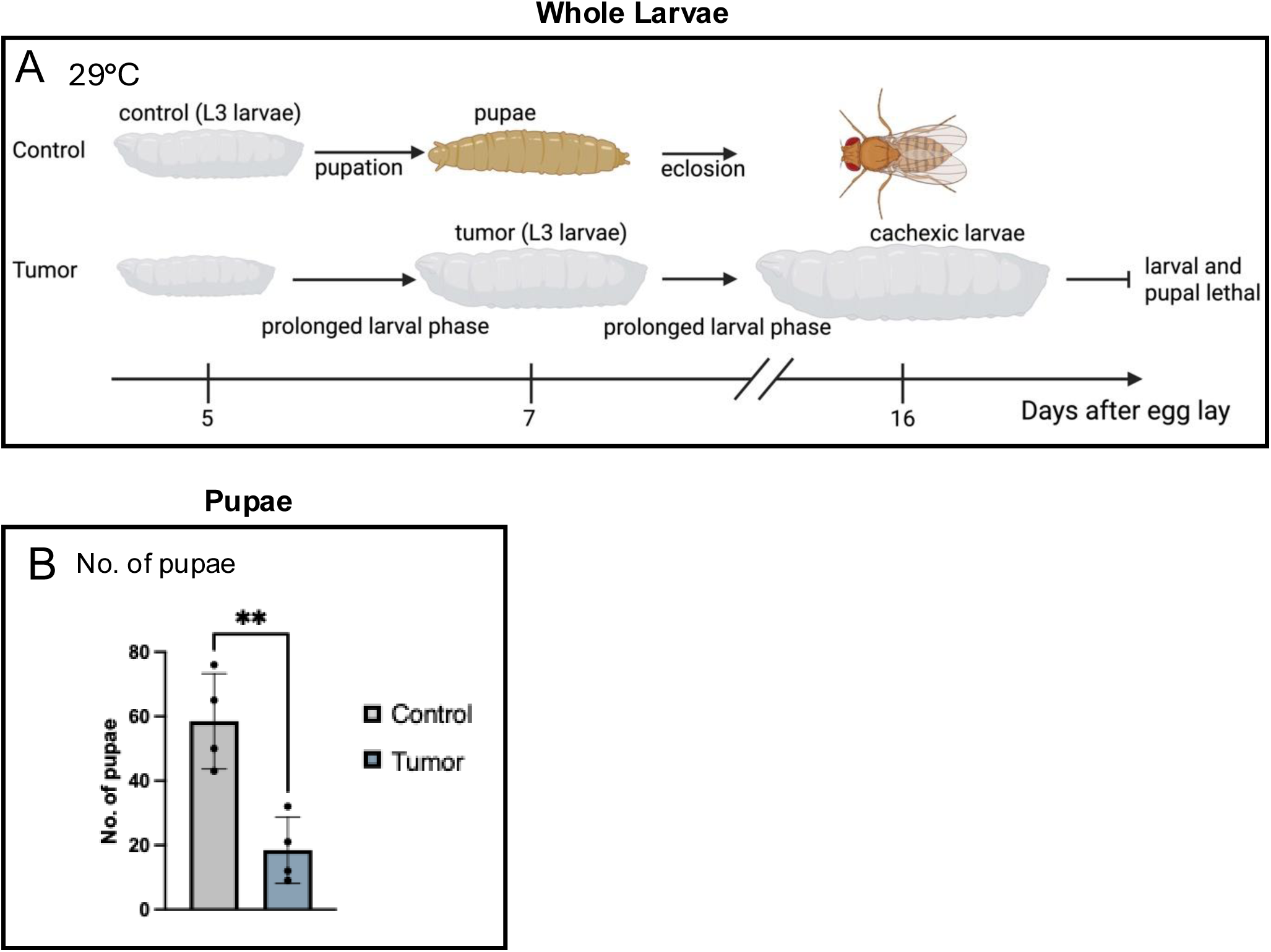
Overexpression of Hipk and Sik-3CA significantly reduces the number of pupae (to fig 1) (A) Schematic diagram depicting development of control and tumor-bearing larvae at 29°C. Quantification of number of pupae of the indicated genotypes. (n=4 biological replicates) ∗∗p = 0.0058 (Welch’s t-test).

**Supplemental Fig. 2.**
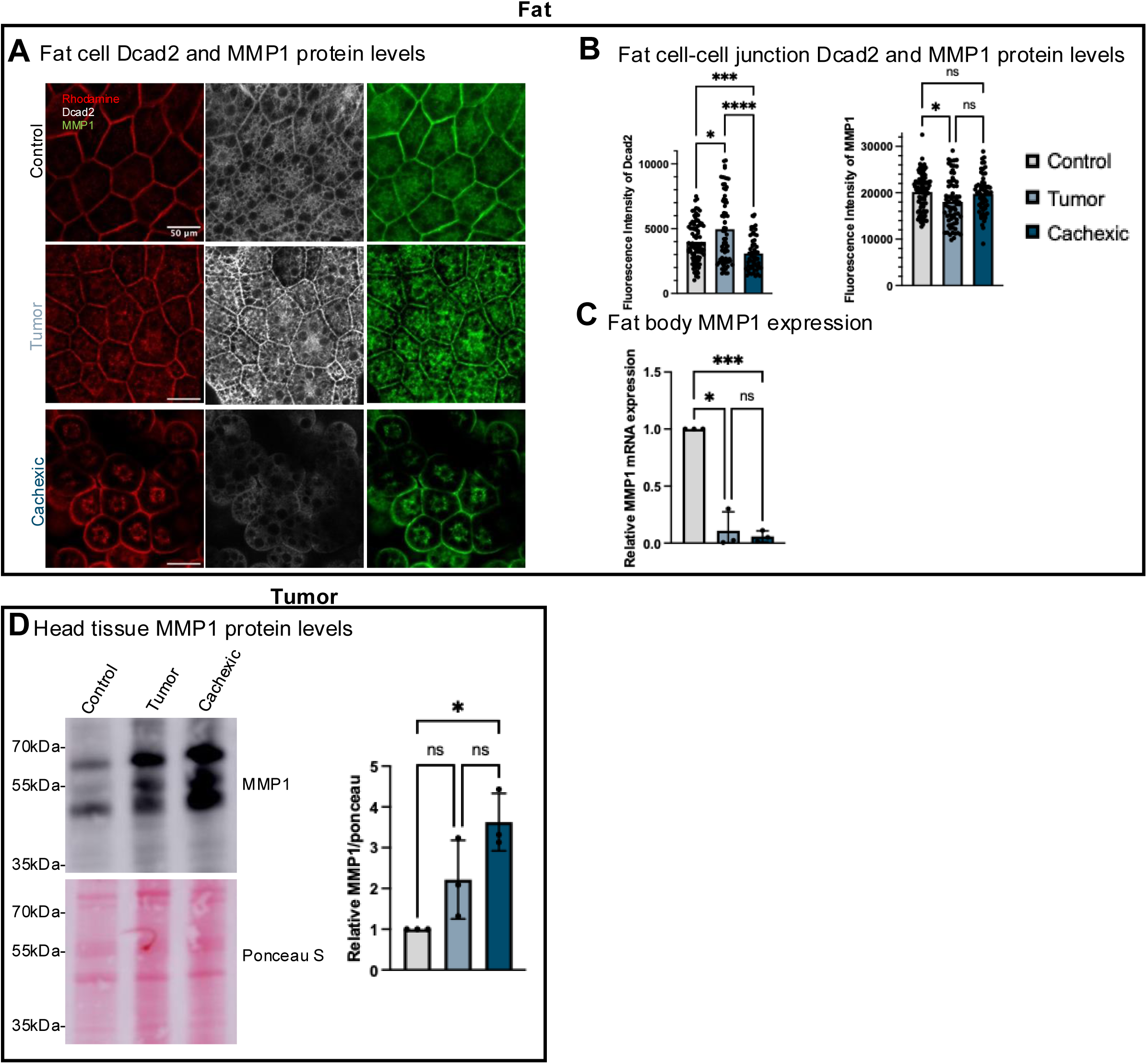
Tumor potentially secretes MMP1 to reduce E-cadherin levels around fat cells causing rounding of fat cells (to fig. 2) (A) Single slice confocal images of F-actin staining (red), Dcad2 (white) and MMP1 (green). (n=27, 21 and 15) (B) Quantification of fluorescence intensity of fat cell-cell junction Dcad2 and MMP1 levels. (n=89, 70 and 58) ns=0.9172 and 0.1061 respectively, ^∗^p = 0.0225, ^∗∗∗^p = 0.0007 and ^∗∗∗∗^p < 0.0001 (Brown-Forsythe and Welch ANOVA test) (C) Fat body MMP1 mRNA levels measured by qPCR (n=5 larvae/data point) *n* = 3 biologically independent experiments ns=0.7735, ^∗^p = 0.0205 and ^∗∗∗^p = 0.0006 (ANOVA Tukey’s multiple comparisons test). (D) Western blot of whole protein extracts of head tissue (including brain, epithelial discs and salivary glands) from third instar larvae and probing for the presence of MMP1 protein. Ponceau S is used as a loading control. *n* = 3 biologically independent experiments (E) Ratio of MMP1 protein level to ponceau S. ns=0.2768 and 0.3485 respectively, and ^∗^p = 0.0418 (ANOVA Tukey’s multiple comparisons test).

**Supplemental Fig. 3.**
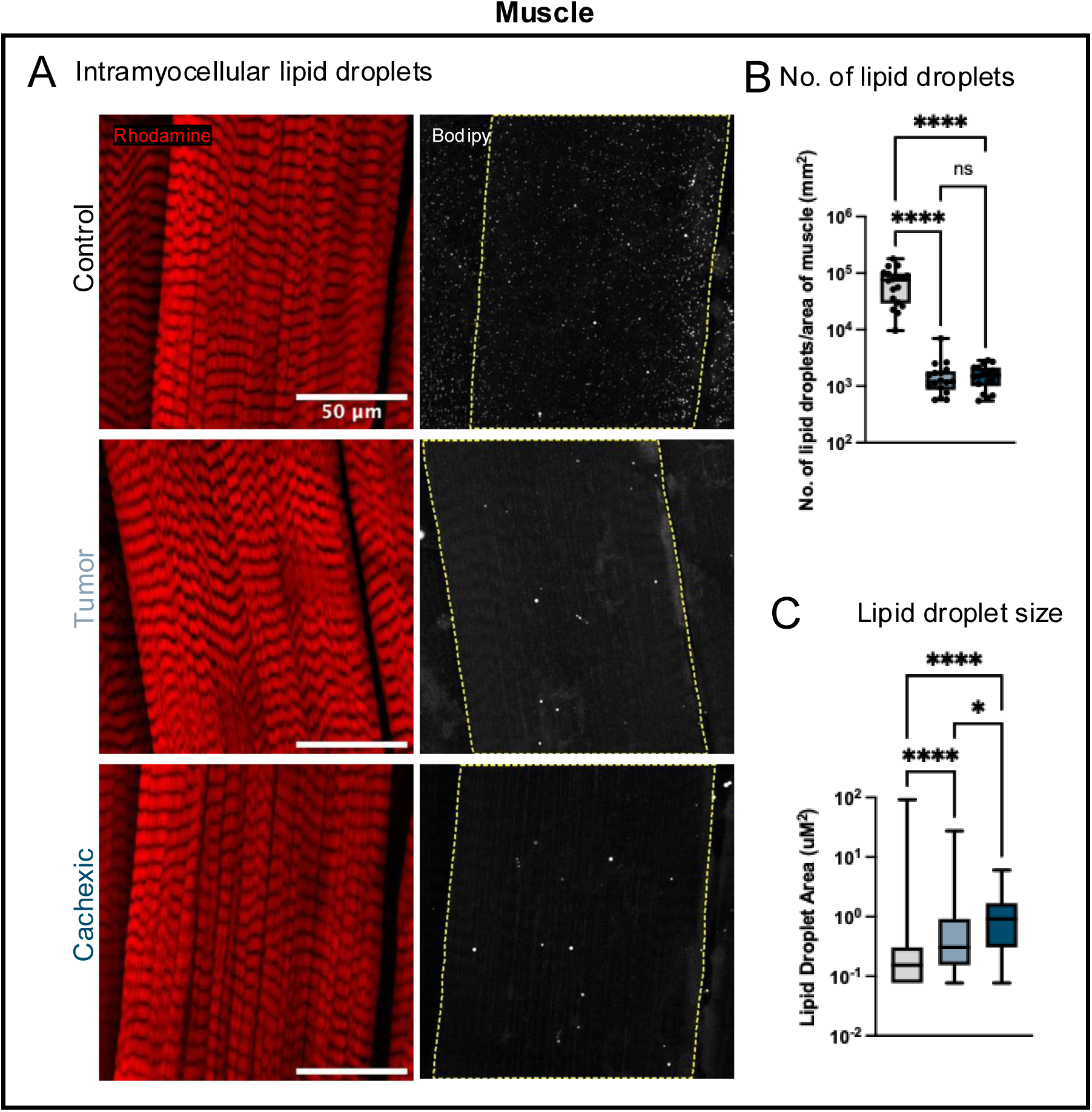
Intramyocellular lipid droplets is significantly reduced in number and increased in size in tumor-bearing and cachexic larvae (to fig 3) (A) Representative longitudinal maximum projection confocal images of of ventral longitudinal VL4 from A2-A4 hemisegments. F-actin (red) and Bodipy (white). (B) Quantification of number of lipid droplet within in each muscle. (n=18, 17 and 18 respectively) ∗∗∗∗p < 0.0001 (Brown-Forsythe and Welch ANOVA test). (C) Quantification of area of lipid droplet within in each muscle. (n=14962, 395 and 417 respectively) ∗p = 0.0263 and ∗∗∗∗p < 0.0001 (Brown-Forsythe and Welch ANOVA test).

**Supplemental Fig. 4.**
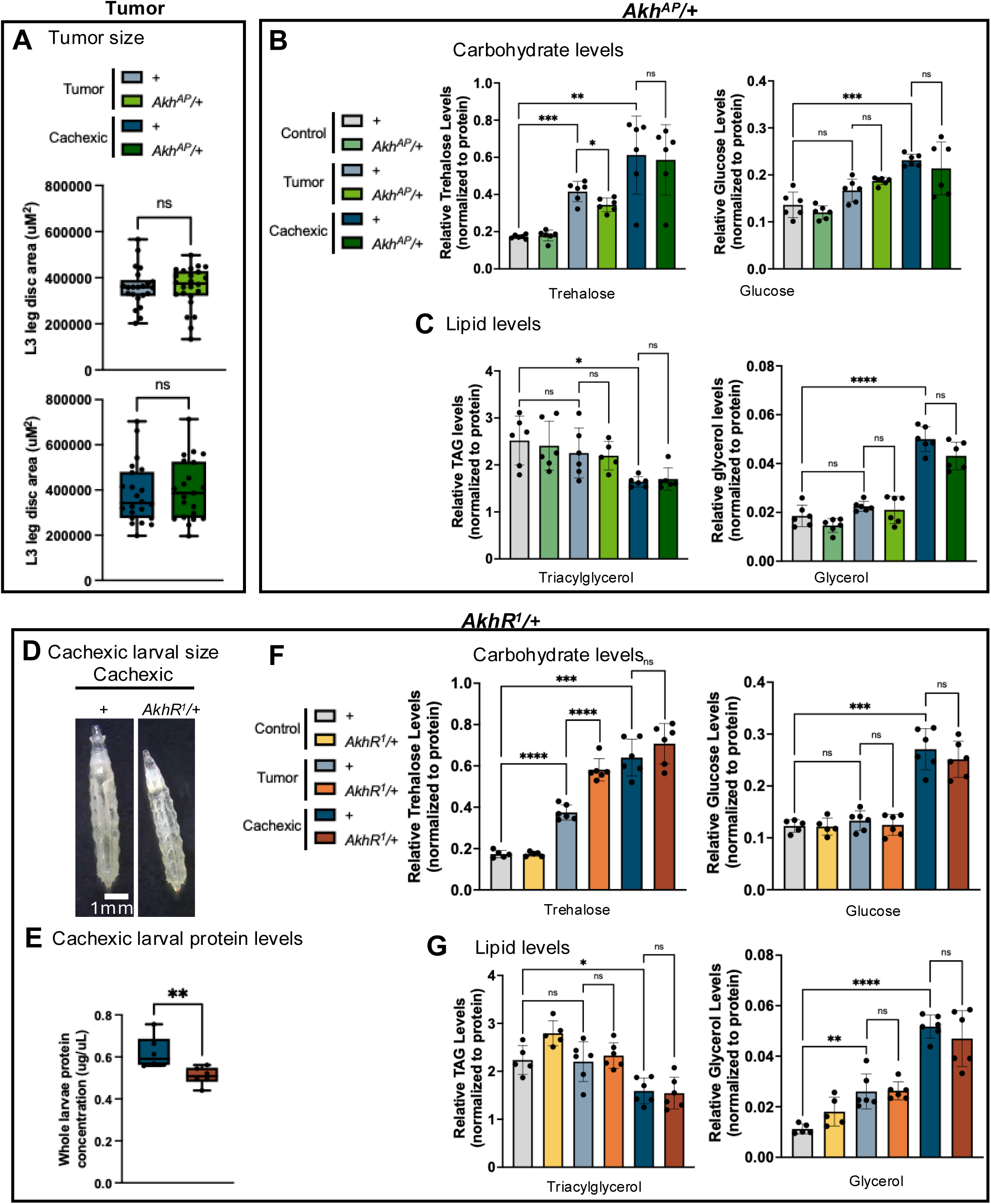
Whole larval partial depletion of Akh can significantly reduce increased trehalose in tumor-bearing larvae but is unable to rescue increased lipolysis and trehalose in cachexic larvae (to fig 5) (A) Quantification of L3 leg disc area for tumor and cachexic larvae (n=22, 24, 22 and 21 respectively) ns = 0.9771 and 0.6753 respectively (Welch’s t test) (B) Trehalose levels in whole larvae normalized to protein levels. (n=5 larvae/data point) ns = 0.826, ^∗^p = 0.0266, ^∗∗^p = 0.007 and ^∗∗∗^p = 0.0002 respectively (Brown-Forsythe and Welch ANOVA test and Welch’s t test). Glucose levels in whole larvae normalized to protein levels. (n=5 larvae/data point) ns = 0.1212, 0.4858 and 0.1045, ^∗∗∗^p = 0.002 respectively (Brown-Forsythe and Welch ANOVA test). (C) Triacylglycerol levels in whole larvae normalized to protein levels. (n=5 larvae/data point) ns=0.313, 0.8305 and 0.6057, ^∗^p = 0.0186 respectively (Brown-Forsythe and Welch ANOVA test). Glycerol levels in whole larvae normalized to protein levels. (n=5 larvae/data point) ns=-.1598, 0.0523, and 0.5639, ^∗∗∗∗^p < 0.0001 (Brown-Forsythe and Welch ANOVA test). (D) Representative images of cachexic larvae (n = 10 and 11 respectively). (E) Protein concentration in whole larvae. (n=5 larvae/data point) ^∗∗^p = 0.0039 (Paired t test). *n* = 6 biologically independent experiments (F) Trehalose levels in whole larvae normalized to protein levels. (n=5 larvae/data point) ns = 0.2402, ∗∗∗p = 0.0001 and ∗∗∗∗p < 0.0001 respectively (Brown-Forsythe and Welch ANOVA test and Welch’s t test). Glucose levels in whole larvae normalized to protein levels. (n=5 larvae/data point) ns = 0.5169, 0.4904 and 0.3855, ^∗∗∗^p = 0.0003 respectively (Brown-Forsythe and Welch ANOVA test). (G) Triacylglycerol levels in whole larvae normalized to protein levels. (n=5 larvae/data point) ns=0.9845, 0.799 and 0.5487, ^∗^p = 0.0113 respectively (Brown-Forsythe and Welch ANOVA test). Glycerol levels in whole larvae normalized to protein levels. (n=5 larvae/data point) ns=0.93, and 0.3657, ^∗∗^p = 0.0046, and ^∗∗∗∗^p < 0.0001 (Brown-Forsythe and Welch ANOVA test).

## Notes

### Competing Interest Statement

The authors have declared no competing interest.

